# Functional succession of actively growing soil microorganisms during rewetting is shaped by precipitation history

**DOI:** 10.1101/2022.06.28.498032

**Authors:** Ella T. Sieradzki, Alex Greenlon, Alexa M. Nicolas, Mary K. Firestone, Jennifer Pett-Ridge, Steven J. Blazewicz, Jillian F. Banfield

## Abstract

Rewetting of seasonally dry soils induces a burst of microbial activity and carbon mineralization that changes nutrient availability and leads to succession. Yet the microbial functions that underpin this succession are not well described. Further, it’s unclear how previous precipitation frames microbial capacities after rewetting and how long these effects persist. We used isotopically-labeled water to rewet seasonally dry annual grassland soil that experienced either mean annual or reduced precipitation during the previous two years, and sampled at five subsequent time points. We used quantitative stable isotope probing (qSIP)-informed genome-resolved metagenomics to identify growing microorganisms, predict their capabilities, and analyze how these traits differed over time and between precipitation treatments. Organisms that grew after wetup were more abundant than non-growing organisms prior to the wet-up, suggesting that traits that initiate succession are pre-defined at the end of the prior plant growing season or via survival over the summer. Fast-growing organisms had fewer carbohydrate active enzyme (CAZy) genes per genome than slower-growing organisms, suggesting that although fast-growers were capable of degrading complex C, they may not specialize in this process. Differential abundance of CAZy genes in growing organisms throughout the succession implies that substrate availability varied with time. In contrast, the abundance of peptidases in growing organisms differed between precipitation treatments, but not over time following wet-up. Before wet-up, the soil organisms’ gene inventories were different between the two precipitation treatments. Surprisingly, this legacy effect waned after just one week. Thus, pre-wetup differences in microbial functional capacity converged shortly after rewetting.

## Introduction

Following the dry season in arid and semi-arid climates, soil rewetting is a key event that triggers microbial community restructuring and substantial carbon turnover [1, 2]. This sudden increase in soil moisture has a disproportionate influence on the annual carbon budget [3] and leads to an immediate release of carbon dioxide, termed the “Birch effect”, which can arise from both abiotic and biotic processes [4–6], with biotic processes dominant in non-carbonate-rich soils [2]. Rewetting causes rapid habitat changes and a large microbial response to the altered soil environment within the span of hours: from survival and dormancy, to rapid osmotic changes and cell lysis, to peak activity [7]. These changes parallel a rapid community succession, with high rates of both growth and mortality that vary by taxonomic group [8]. While the identities of the bacterial and archaeal taxonomic groups that respond to rewetting have been described, the physiological traits that underpin organism interactions and enable rapid microbial growth and turnover following wet-up are poorly understood.

Prior research has investigated the organic resources that drive the pulse of biogeochemical processing in Mediterranean-climate grassland soils following the first autumn rain event. In addition to dry plant detritus, labile carbon and nitrogen accumulate over the summer dry season [9], largely due to low microbial metabolic rates and reduced microbial uptake [10].

Sustained activity of secreted extracellular enzymes [10, 11], osmotic shock [12] and secretion of compatible solutes also contribute to the buildup of labile C and N pools [13–15]. Soil rewetting is generally followed by high degradation rates of these substrates [16], supporting initially fast-growing organisms [17]. The existing paradigm suggests these fast growers rely on labile resources. However, we know little about the metabolic characteristics of the initially fast-growing organisms [8] which are significant contributors to the Birch effect.

In Mediterranean systems, climate change is predicted to cause increased water stress [18] due to decreased rainfall, extension of the dry season and increased frequency of drying/rewetting cycles, leading to more CO_2_ pulses [19, 20]. It has been noted that there is less immediately respirable carbon when the dry season is prolonged [10]. We hypothesized that reduced precipitation in the preceding growing season could affect the response of soil microbial communities to rewetting following the long dry summer. Specifically, we predicted that differences in prior year water input would lead to differences in the biogeochemical toolkit of organisms that grow following wet-up, and that these differences would persist over time.

Here, we used stable isotope-informed genome-resolved metagenomics to identify and analyze the genomes of organisms that become active in the week following soil wet-up. From these genomes, we predicted the functional capacities of growing organisms and investigated how they changed over time and due to differences in rainfall history. Our results reveal distinct gene inventories of fast-responding organisms after wet-up in soils that had experienced two distinct rainfall treatments. We document shifts in functional capacities over time, consistent with changes in carbon compound resources, and convergence in the capacities of growing organisms after approximately one week.

## Results

Surface soil samples from a Mediterranean annual grassland with a reduced vs. mean precipitation experiment were collected in the late summer of 2018, prior to the first autumn rainfall event, and used for a ^18^O-water quantitative stable isotope probing (qSIP) wet-up experiment. DNA was extracted pre-wetup (0h) from soil that experiences mean or reduced precipitation, at 4 time points after water addition from the mean precipitation treatment samples (24h, 48h, 72h and 168h) and 2 time points from the reduced precipitation treatment. The DNA was density fractionated, and 5-7 Gbp of Illumina sequences were acquired per density fraction in addition to 20-28 Gbp from unfractionated DNA per sample. 338 dereplicated metagenome assembled genomes (MAGs) were generated from this dataset and combined with 168 MAGs from a previous metagenomic study from a nearby field site sampled during winter [21]. These 503 genomes representing 11 phyla and 23 classes of bacteria and archaea were at least 75% complete. Three additional Firmicutes MAGs, the only representatives of this phylum, were of low quality and therefore used only for abundance and growth analyses. Of the total pool of MAGs, 199 had a breadth (covered fraction) of at least 50% in at least one sample from this study. Failure to detect the remainder of the MAGs (304/503) with coverage above this threshold may be because coverage of the genomes in any individual sample was too low to enable assembly (thus they were only reconstructed via co-assembly of data from multiple samples). Short reads mapped to the MAGs made up an average of 12% of total reads per unfractionated sample and 24% of reads from the density fraction samples. This level of assignment of reads to genomes is expected because the majority of DNA from soil originates from very low abundance organisms [22, 23].

Atom fraction excess (AFE), a metric of growth-based activity, was calculated per MAG based on isopycnic fractionation (fractionation by ultracentrifugation based on changes in DNA density resulting from heavy isotope incorporation, in this case ^18^O). 79 MAGs had statistically significant isotopic enrichment in at least one time point of one watering regime, as indicated by AFE lower confidence interval limit > 0. We refer to this subset of organisms as growing, since isotope enrichment of DNA requires replication. Hereafter, we distinguish between analyses that include all 506 organisms and analyses of the 199 organisms that were detected by read mapping, of which 79 were growing (sup. table S1).

We used 16S rRNA based quantitative stable isotope probing (qSIP) to characterize changes in microbial community structure and determine differences in post wet-up mortality in soils that had experienced mean precipitation or reduced precipitation during the previous growing season. We also measured total soil respiration (CO_2_) and new plant-biomass derived respiration (i.e., decomposition of carbon fixed during the preceding growing season labeled with ^13^C). While total soil respiration was not significantly different between the precipitation treatments (fig. 1A), the microbial mortality rate and respiration of newly-fixed plant material were lower in the reduced precipitation treatment (fig 1B, C, p=0.08). The cumulative microbial mortality rate was three orders of magnitude higher in the mean precipitation treatment.

**Figure 1:**
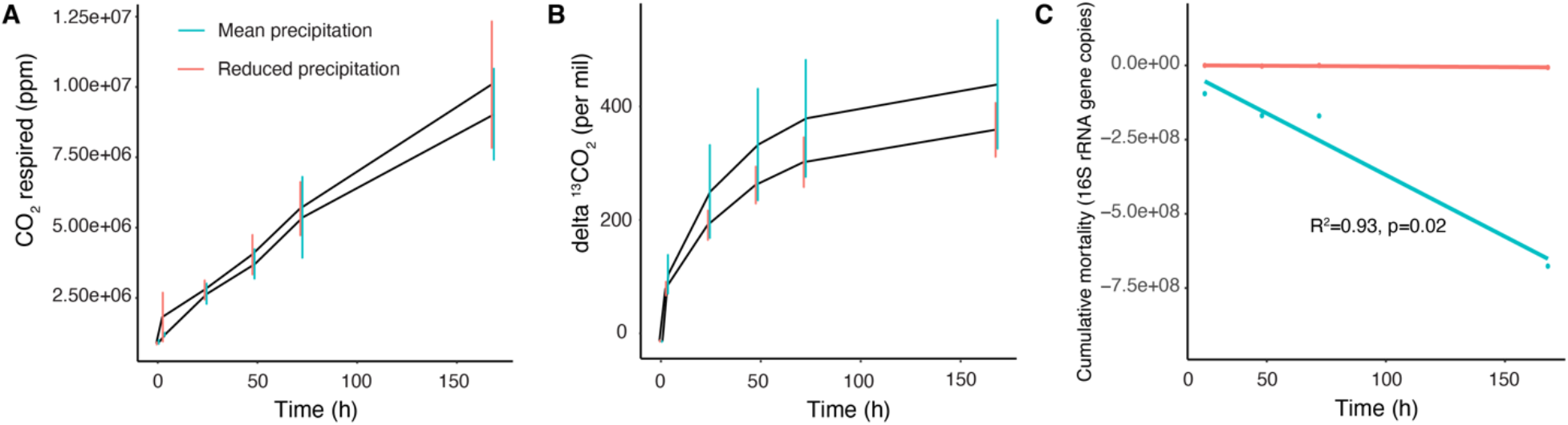
Effects of a 1 week wet-up experiment and two precipitation treatments in annual grassland soils from Hopland, CA: (A) total soil respiration (N=3 per treatment), (B) ^13^CO_2_ respiration of new plant material (N=3 per treatment), (C) cumulative qSIP-based microbial mortality rate, starting at 24h. The linear regression of mortality in the mean precipitation treatment explained 93% of variability with p=0.02. Error bars reflect standard deviation.

To test whether growing microorganisms were more abundant or became more abundant compared to detected but non-growing organisms we calculated the abundance (measured as reads per kilo million, RPKM) of both growing (N=79) and non-growing (N=121) organisms at each post-wetup time point and in both treatments (fig. 2). Growing microorganisms were significantly more abundant than non-growing organisms at time 0 for both treatments (p<0.001). Abundance of growing or non-growing organisms did not vary between precipitation treatments. There was no statistically significant change in mean abundance over the following week for growing or non-growing microorganisms. With that, the variance in abundance of growing and non-growing organisms in the reduced precipitation treatment decreased significantly over the first 48 hours (fig. 2 see asterisks), indicating that the abundance of the rarer growing organisms increased.

**Figure 2:**
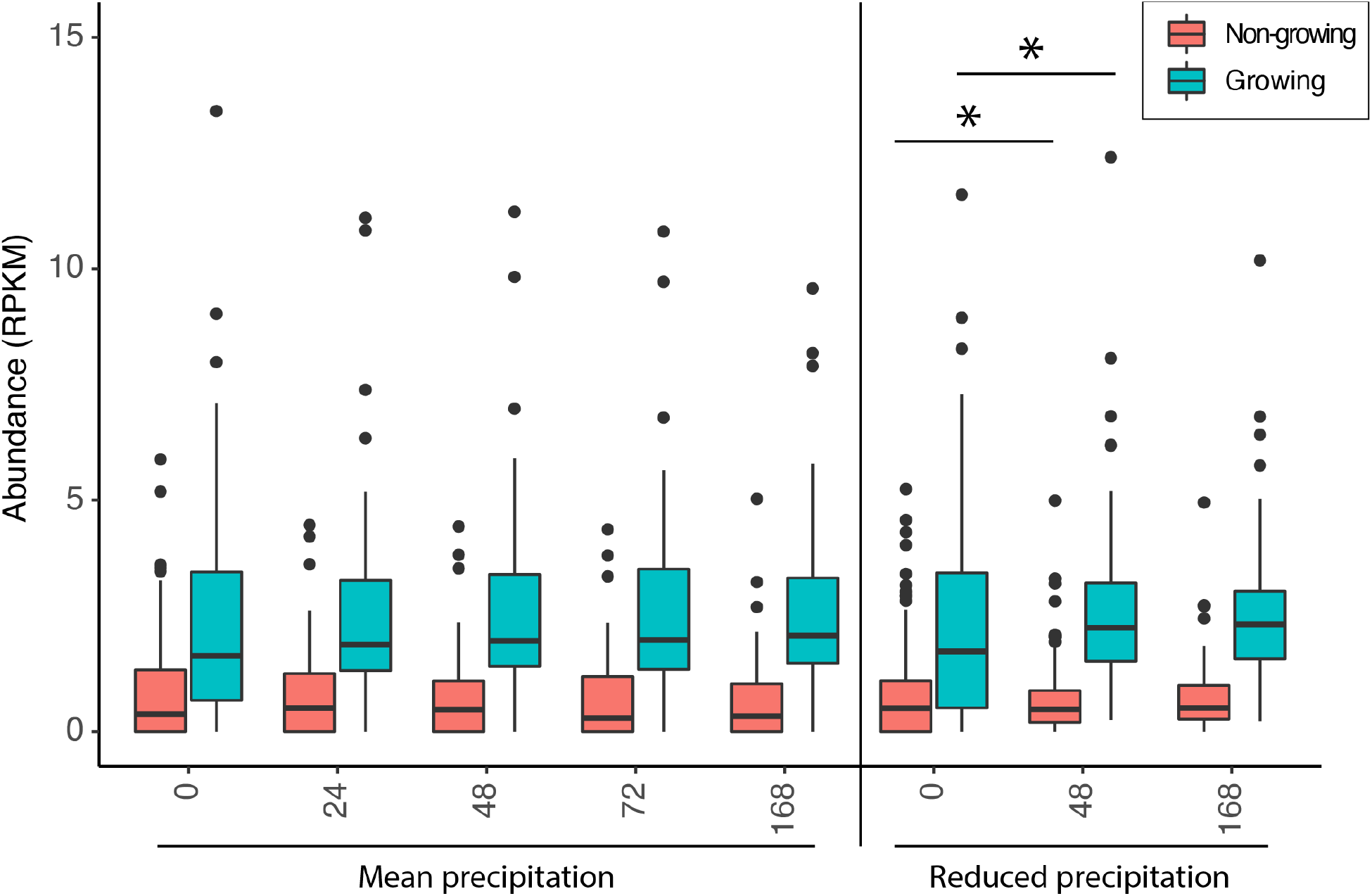
Abundance (RPKM, reads per kilo million) of actively growing versus non-growing bacteria and archaea after a wet-up of dry grassland soils that had experienced mean or reduced precipitation in the preceding growing season. Growth was determined by quantitative stable isotope probing via ^18^O-water incorporation. All differences between growing organisms (N=79) and not-growing but detected organisms (N=120) are statistically significant (p<0.001); within these groups, there are not statistically significant differences with time or precipitation treatment. Boxes represent 75% of that data, whiskers indicate 90%, outliers are denoted by dots. The differences in variance (box length) between 0h and 48h in the reduced precipitation treatment are significant in both growing and not growing organisms (p<0.05).

To test the relationship between abundance and growth, we assessed whether abundant growing organisms were also growing rapidly (AFE > 0.2) by comparing abundances and AFE values of growing microorganisms (N=79) at each time point (fig. 3). Less abundant organisms tended to be fast growing (e.g., Alphaproteobacteria and Gammaproteobacteria) whereas the more abundant organisms were slower growing (e.g., Actinobacteria) over the week post-wetup (linear regression, p<0.001, R^2^=0.1). Significant growth was also detected in Verrucomicrobiae, Gemmatimonadetes, Chloroflexi, Myxococcia, Acidobacteria and the ammonia oxidizing archaea class Nitrososphaeria (fig. 3).

**Figure 3:**
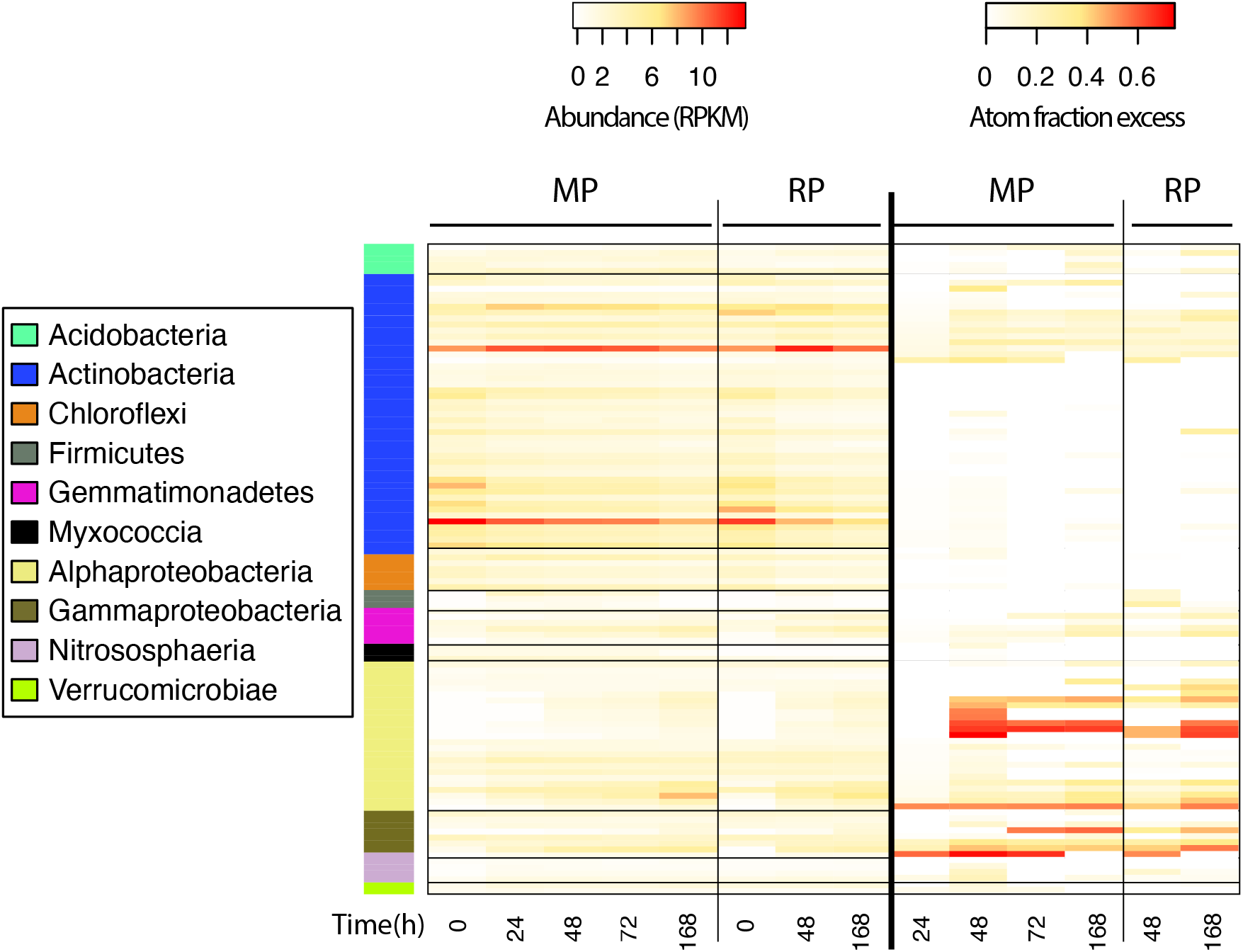
Abundance, enrichment and taxonomic affiliation of 79 taxa that were growing (atom fraction excess confidence interval > 0) after a grassland soil wet-up with ^18^O enriched water, illustrating that highly abundant organisms in both mean and reduced precipitation treatments were slow growing, and vice versa. Each row represents one taxon, and each column represents one time point. The heat map on the left shows abundance normalized to sample size and MAG length (RPKM) and the heat map on the right shows ^18^O enrichment (AFE). The rows are ordered by class and enrichment within class, and the left bar represents taxonomy. MP = Mean precipitation, RP = reduced precipitation. Negative AFE values were considered undetectable and converted to zero.

Most growing organisms were isotopically enriched at multiple consecutive time points, with 63 out of 70 and 30 out of 40 in the mean precipitation and reduced precipitation soils respectively (fig. 3). To distinguish boom and bust growth dynamics compared to persistent enrichment on an organism-by-organisms basis, we plotted the number of growing organisms only at one time point vs. in multiple time points (fig. 4). A bloom of 28 newly growing organisms occurred at 48h in the mean precipitation treatment, but the majority of these organisms did not continue growing later (see leftmost column in fig. 4), representing boom and bust dynamics. In contrast, for 10 organisms, mostly Proteobacteria, we detected growth in all time points in both treatments (see rightmost column in fig. 4).

**Figure 4:**
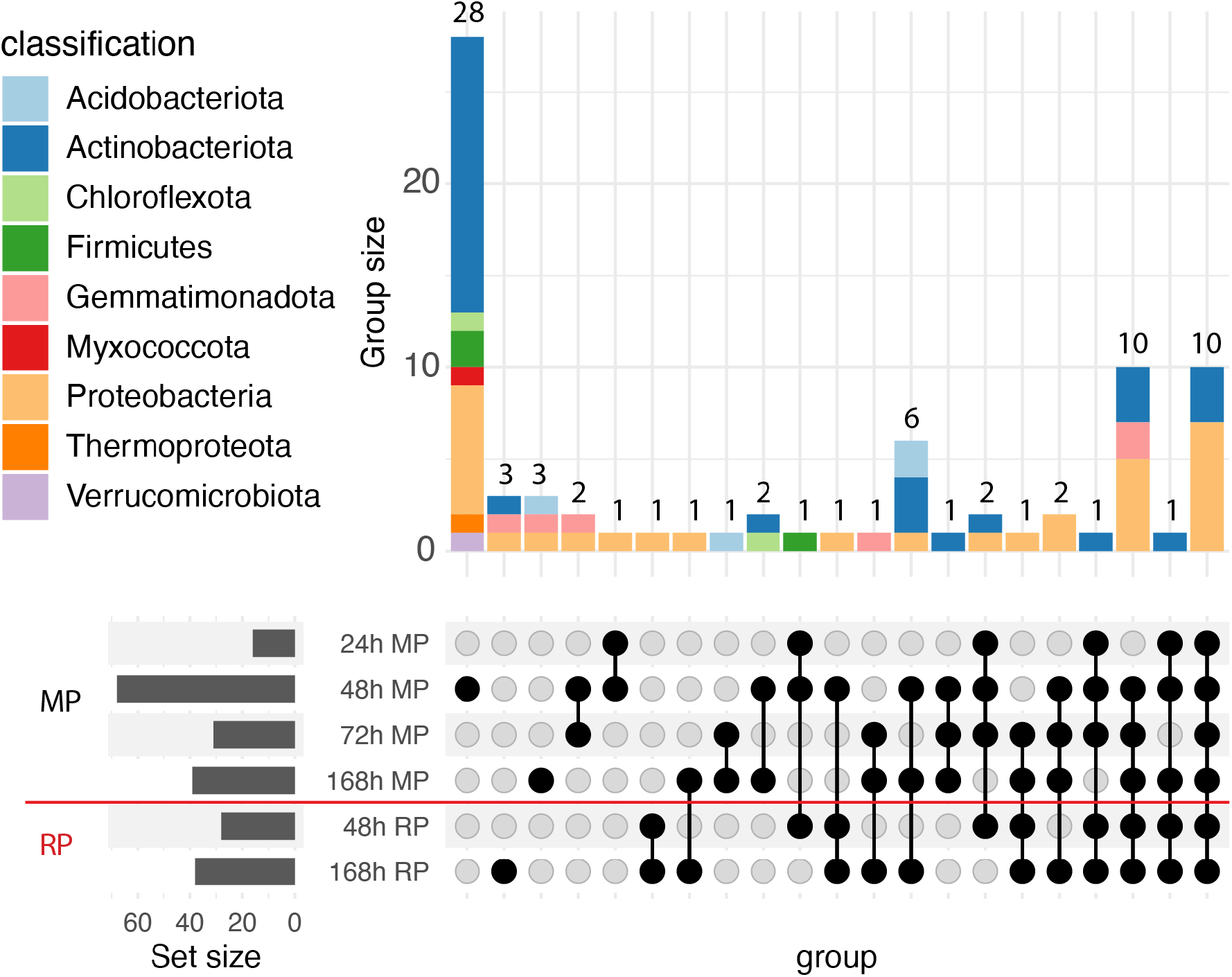
Growth temporal patterns in microbes during the week following wetup of Mediterranean soil that experienced mean or reduced precipitation during the two previous growing seasons. Each row represents a time point in a single precipitation treatment. A single dot per column represents organisms growing only in that time point, whereas linked dots represent organisms growing in both time points/treatments. Each column, denoted by either a single dot or linked dots, represents a group (e.g., organisms that grow at 48h, 72h and 168h in the mean precipitation treatment). Each bar shows how many organisms were growing within a specific group (group size), as well as their taxonomic distribution. In both mean (MP) and reduced precipitation (RP) treatments, when grouped using growth patterns over time (black dots), it is apparent that many organisms grew in just one time point, others displayed a longer boom before bust growth pattern and some grew continuously.

To probe the extent to which changes in genetic repertoires might underpin shifts in organism abundances we compared the abundances of annotated genes in growing organisms in all samples. The purpose of this gene-centric approach was to identify genes common in growing organisms, as opposed to examining each genome of a growing organism and searching for common genes. We interpret a higher number of genes encoding a trait in growing organisms as an indication of a higher capacity of the community to perform that function. Out of 275,788 annotated genes from growing organisms (N=79), the abundance of 30,175 genes changed significantly (padj < 0.05), either between consecutive time points or between treatments at the same time point. In the mean precipitation treatment, the number of genes changing in abundance between consecutive time intervals varied by two to three orders of magnitude. This was not apparent in the reduced precipitation treatments (fig. 5A). However, at time 0h, when comparing the mean vs. reduced precipitation treatments, 11,149 genes were significantly more abundant in one of the treatments (fig. 5B). Most genes at this time point were more abundant in the mean precipitation treatment (sup. fig. 1). This effect waned over time and disappeared completely at 168h (fig. 5A). Ordination of gene abundance in growing organisms revealed progressive temporal changes in community gene abundance in both precipitation treatments (fig. 5C).

**Figure 5:**
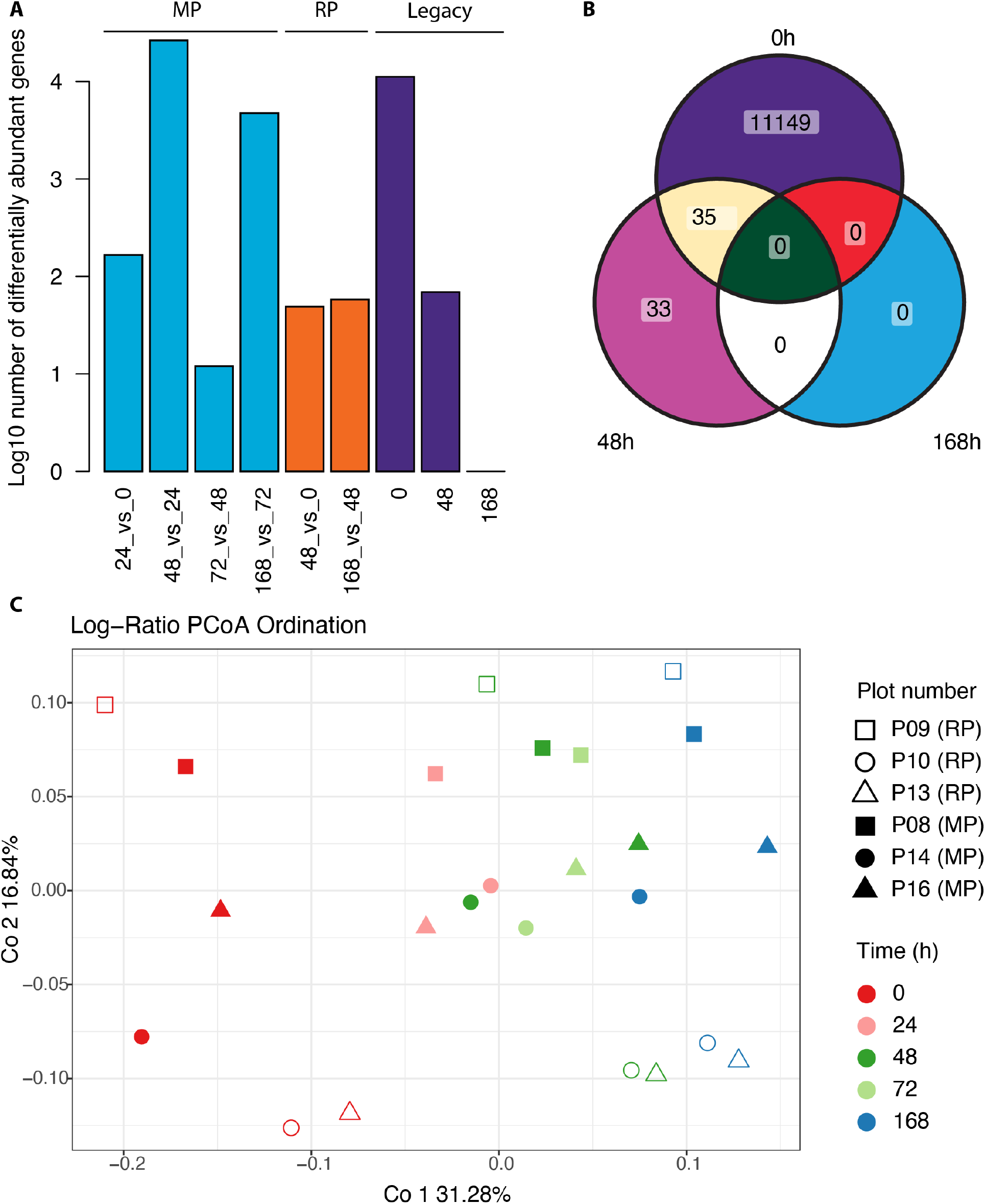
Changes in gene abundance over time and between precipitation treatments reflect changes in abundance of growing organisms after soil wetup. (**A**) Number of significantly (p<0.05) differentially abundant open reading frames (ORFs) at each time point compared to the previous time point (e.g., 48h vs. 24h). Three main comparisons are represented by distinct colors: Mean precipitation treatment (**MP, light blue**), Reduced precipitation treatment (**RP, orange**) and comparison between the two at the same time point (**legacy**, **dark purple**). Note that the y axis scale is logarithmic. (**B**) Venn diagram of differentially abundant genes between MP and RP precipitation at the same time point (legacy effect). (**C**) Principal coordinates analysis of log-ratio transformed abundance of genes from growing organisms normalized to sequencing depth (N=79). Shapes represent the location of the field plot from which the metagenome was sequenced: the upper row (plots 8 and 9) or the lower row (plots 10, 13, 14 and 16; see methods). Full shapes denote MP, and empty shapes RP. The first two components explain 48% of the variance.

Dynamics of differentially abundant genes differed for mean and reduced precipitation treatments. In the mean precipitation treatment, the number of differentially abundant genes mostly decreased over the first day, became more abundant over the following day, decreased over the third day and some were more and others less abundant by the end of the week (sup. fig. 1). The reduced precipitation treatment showed a distinct pattern: all differentially abundant genes were more abundant after one week (sup. fig. 1, panel RP 168h vs. 48h). Comparing time points between treatments indicates that most genes were initially (0h) more abundant in the mean precipitation compared to the reduced treatment (sup. fig. 1).

Within a given treatment, we tested whether the genes that changed in abundance between consecutive time intervals were the same genes. In the mean precipitation treatment, 88% of differentially abundant genes were unique to the 48h time point (sup. fig. 2A). In the reduced precipitation treatment, 99/182 differentially abundant genes were shared between 48h and 168h (sup. fig. 2B). Almost all the genes (472/510) that differed at time 48h also differed at 0h (sup. fig. 2C).

To search for evidence of factors that may have driven changes in growth behavior over time, we evaluated inventories of genes for degradation of carbohydrates, including plant material, fungal material, and other polysaccharides. These functions are performed by carbohydrate active enzymes (CAZy). The difference in abundance patterns of CAZy genes between growing organisms (N=79) and detected but not growing organisms (N=120), i.e., which CAZy genes were more abundant than others, was significant (Wilcoxon rank sum test, p<0.001). Thus, organisms that grew during wet-up possessed a different set of CAZy pathways compared to the rest of the bacterial and archaeal community. We looked for a correlation between the number of CAZy genes per genome and atom fraction excess, representing growth. We found that fast growing organisms (AFE>0.2, based on sup. fig. S3) had significantly fewer CAZy genes per genome (average 33 genes per genome), compared to an average of 41 genes per genome in slow growers (sup. fig. 3; one sided Student’s t-test, p=0.02, t=-2). To verify that this difference was not due to lower genome completeness of fast-growing organisms, we compared their genomes completeness values to those of slow-growers and found no significant difference (Student’s t-test, p=0.66).

Because complex carbohydrates are an important carbon source in soil not just during wetup but throughout the year, we compared CAZy pathways found in growing organisms to CAZy repertoires of the whole genomically sampled community, defined as all organisms detected in the wetup and during winter (N=503)[21]. We found genes coding for the degradation of rhamnose, pectin, beta mannan, mixed linkage glucans, fucose, chitin, crystalline cellulose more highly represented in growing organisms than in the total set of organisms (sup. fig. 4).

The total abundance of certain CAZy pathways, especially those related to plant material degradation, changed over time in growing organisms (fig. 6). For example, degradation of amorphous cellulose (C) became relatively more abundant between 24h and 48h. In contrast, xyloglucan (X) degradation became relatively less abundant between 24h and 72h. Based on the percent of growing organisms containing specific CAZy pathways, we observed a functional succession mainly of plant material degrading enzymes. These changes were statistically significant between time points in the mean precipitation treatment, as well as between treatments at 48h, but differences between treatments were not measurable 168h after wetup.

**Figure 6:**
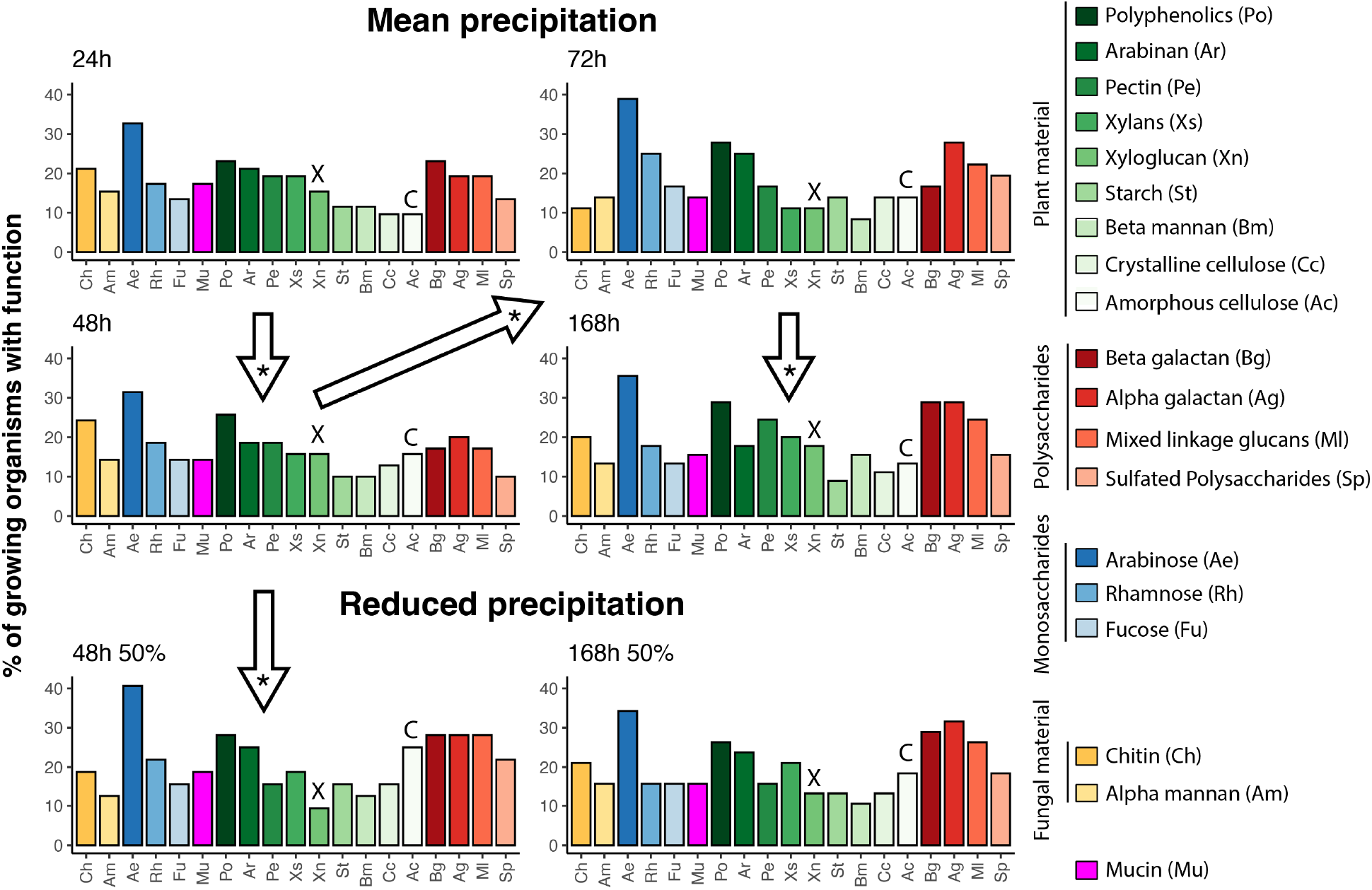
Percent of the organisms that were growing during the week following soil wetup at each time point and contained genes coding for carbohydrate degradation pathways in order of time and precipitation treatment. Arrows indicate significant changes to the entire profile determined by Wilcoxon rank sum test (Benjamini-Hochberg correction for multiple comparisons p<0.05). Xyloglucan degradation is marked by an X and Amorphous cellulose by a C above the respective bars.

We also investigated nitrogen acquisition and cycling pathways in growing organisms. Nitrogen in soil exists mostly in the form of large, complex molecules such as proteins, chitin and nucleic acids, but it may also be present as ammonium and other inorganic nitrogen forms [24]. Some inorganic forms of nitrogen can be used as an energy source, such as oxidation of ammonium in nitrification. To investigate what forms of nitrogen growing organisms were taking up or transforming during wet-up, we compared the percentage of growing organisms that contained various nitrogen related genes to the genetic repertoire of the year-round whole community. We found enrichment of genes related to ammonium assimilation and ammonium transporters in growing organisms (Student’s paired t-test, p=0.06, fig. 7A), but no enrichment of organic nitrogen acquisition genes (Student’s paired t-test, p=0.33, fig. 7B) except for peptidases (Student’s paired t-test, p=0.1, fig. 7C). Growing organisms contained more peptidases per genome that belonged mainly to three catalytic groups: metallopeptidases, serine peptidases and cysteine peptidases (fig. 7C). While the number of peptidases per genome in growing organisms did not change over time, there was a significant difference (p<0.05) between this number in the mean precipitation treatment at times 48 h and 72 h compared to the reduced precipitation treatment at 48 h.

**Figure 7:**
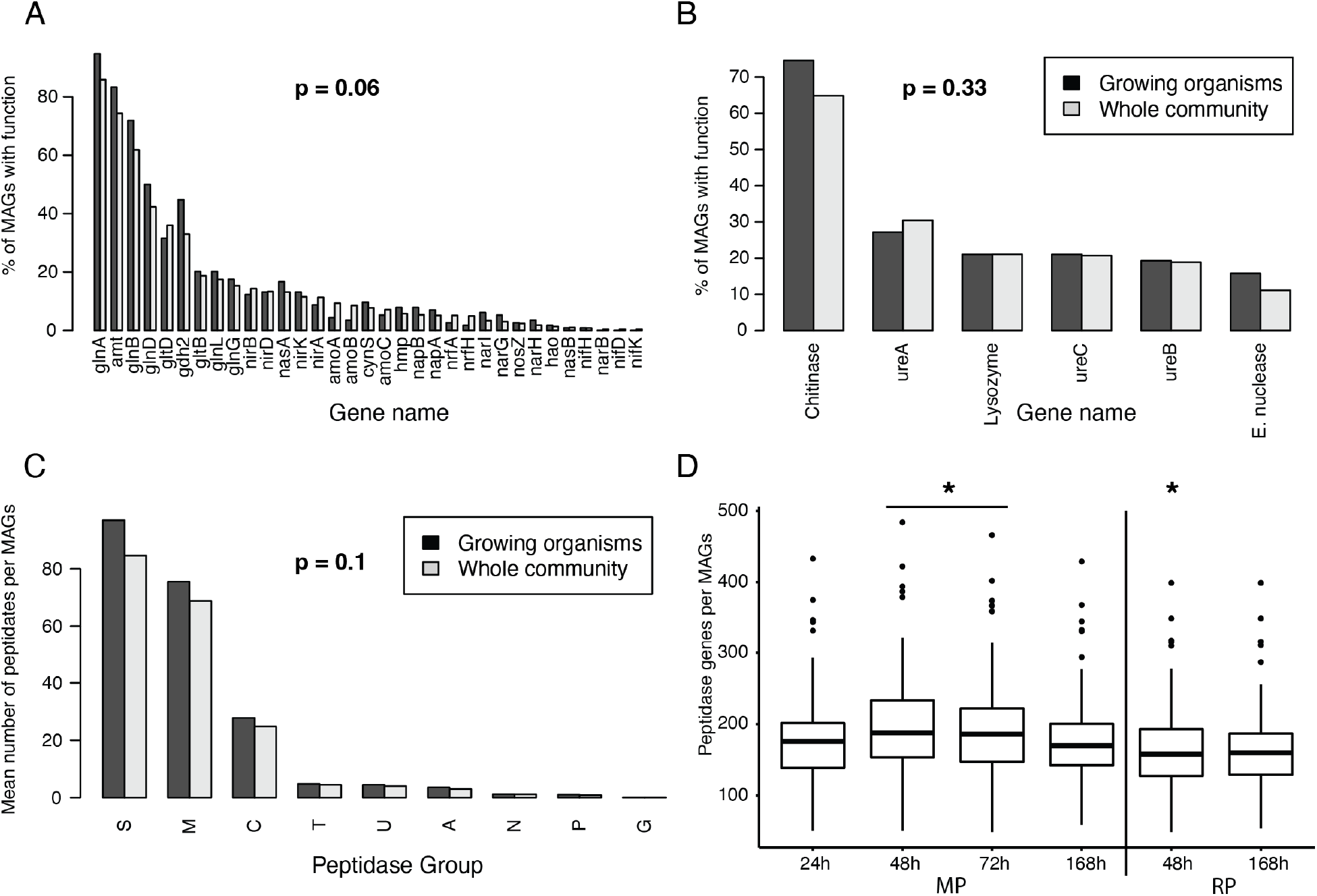
Presence of nitrogen acquisition and cycling traits in growing organisms during the week following soil wetup. Comparison of N acquisition/cycling genes of (A) inorganic N transformations, (B) organic N transformations and (C) catalytic groups of peptidases in growing organisms vs. the whole community. P-values based on Wilcoxon rank sum test are displayed on each panel. Peptidase catalytic groups: S=serine, M=metallo, C=cysteine, T=threonine, U=unknown catalyst, A=aspartic acid, N=asparagine, =mixed, G=glutamic acid. (D) Number of peptidases per organism in growing organisms did not change over time, but the number is significantly (p<0.05) lower at 48h in reduced precipitation compared to 48h and 72h at mean annual precipitation, indicated by asterisks. MP = Mean precipitation, RP = reduced precipitation

Two of the main strategies in which soil organisms can deal with dry conditions are osmolyte biosynthesis and production of extracellular polysaccharides (EPS) [25]. We searched for genes coding for biosynthesis of the osmolyte trehalose and for EPS synthesis and export. We found significantly more EPS export genes in growing organisms compared to the whole community (Student’s paired t-test, p=0.003; fig. 8A) but no significant differences in trehalose synthesis (data not shown). We observed no effect of rainfall treatment or temporal changes in the percent of growing organisms that possess EPS export genes (fig. 8B).

**Figure 8:**
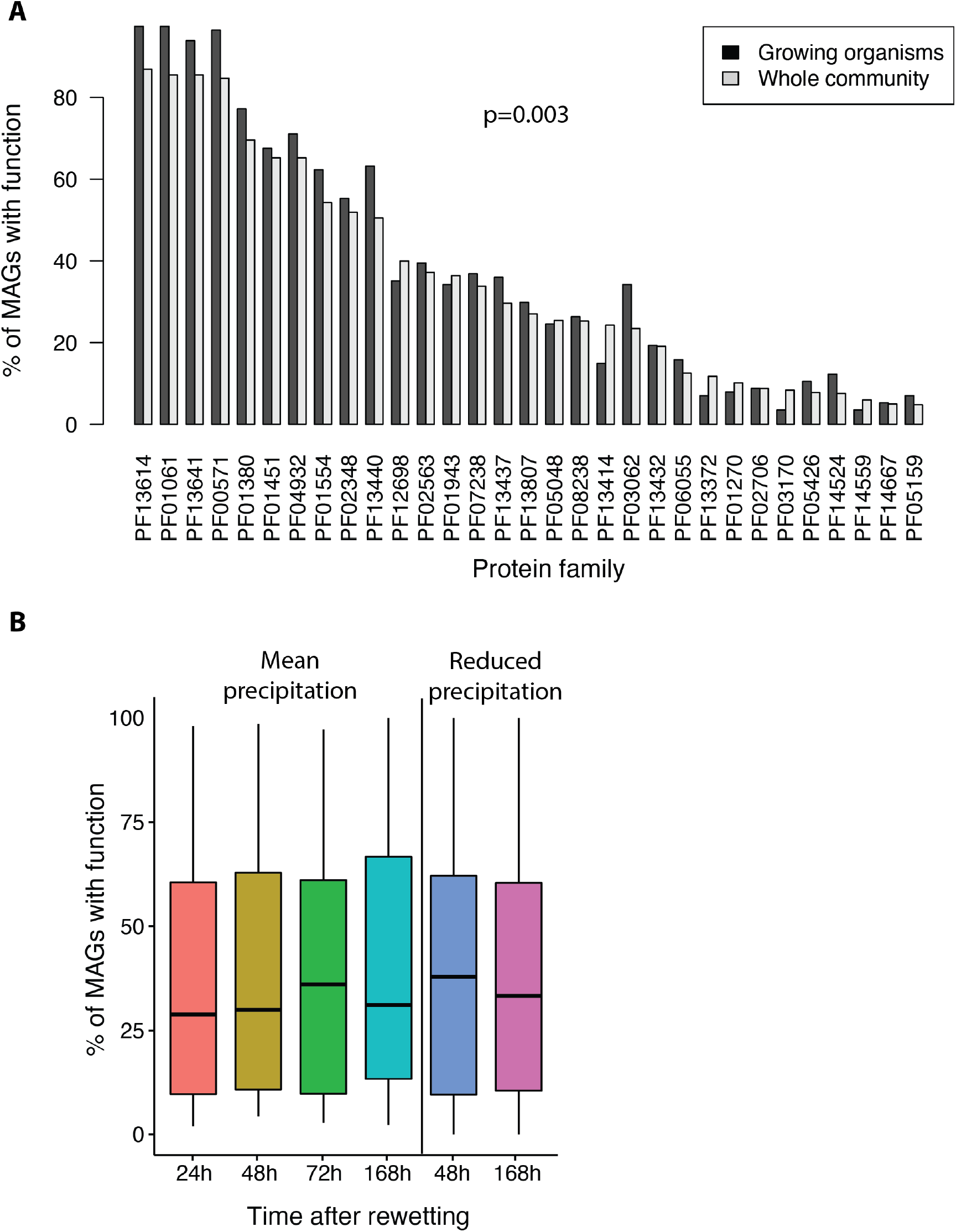
Presence of EPS export genes in metagenome assembled genomes (MAGs) of soil organisms during the week after soil wetup in (A) growing organisms vs. the whole community (Student’s paired t-test, p=0.003), and (B) in growing organisms over time (ANOVA, p=0.99).

## Discussion

We investigated how the history of reduced rainfall impacts the response of soil microbial communities to wet up and the processes by which they acquire nutrients to support growth. While climate change and its associated periodicity of rainfall and longer droughted periods affecting soil moisture are predicted to reduce grassland soil microbial diversity [26], it is the active organisms that determine ecosystem function, therefore we focused on those through stable isotope probing. Integrated results from stable isotope informed genome-resolved metagenomics, gene abundance, and gene function information enabled characterization of the traits of growing organisms that may have contributed to their proliferation during the week following rewetting. Our findings provide a mechanistic foundation for understanding of bulk rate measurements of soil biogeochemical processes following wetup (e.g., respiration).

As conditions following wetup are quite distinct from those during the summer dry period, we expected that organisms that were abundant during dry down would not be those that grow immediately following wet up. However, in the mean precipitation treatment, the organisms that grew over the week following wetup were a subset of those present in high abundance prior to water addition (t0). Upon wetup, growth was almost certainly primed by newly available labile compounds released by cell death due to osmolysis [17, 25]. Cell lysates may also have been available following desiccation during dry down. Thus, growing organisms may simply be those that utilize compounds released upon cell death rapidly.

Organisms that grew following wetup were initially more abundant. It may be that they were abundant immediately prior to dry down, or they survived the dry season better than others, or both. We observed a higher prevalence of genes coding for synthesis and export of extracellular polysaccharides in growing organisms compared to the whole community implying that growing organisms may be better poised to survive dry soil conditions. Thus, in combination, we suggest that organisms that were abundant at t0 and grew following wetup were those capable of surviving summer dry conditions and that can grow rapidly on cell lysates. Moreover, organisms that grow may also rely on more complex carbon sources but be primed by labile nitrogen, such as ammonium and amino acids.

Of the organisms that grew, the fastest growers over the following week were the bacteria that were not the most abundant organisms. Why this is the case remains unclear, but this pattern has been observed previously in grassland soil during the winter [27]. It is possible that organisms that rapidly respond to wetup rely on resources that are less available during most of the year, such as labile carbon and inorganic nitrogen. In support of that, genomes of fast growers contained less carbohydrate degradation genes than those of slow growers, implying a limited toolkit for accessing complex carbon, which is the main carbon source in soil during the rest of the year [28]. A similar pattern of rare and fast-growing bacteria was also identified in a marine environment [29], in which short-lived resource availability was also suggested as a potential mechanism.

Notably, the organisms that were least abundant prior to wetup exhibited no growth at all following water addition. We suggest that these organisms are part of the low abundance microbial “seed bank” and proliferate under conditions that differ from those studied here. Seed banks are a common strategy of microbial communities to overcome unfavorable conditions and likely underlie successional processes [30]. However, we acknowledge that we likely cannot detect growth of all rare organisms due to the noise inherent in qSIP calculations.

We tracked the changes in gene abundance over the week following wetup and noted that gene abundances changed dramatically and over daily timescales. The observation that genes in growing organisms are almost all becoming less abundant (i.e, regardless of the function of the gene) over the first 24 hr is likely attributed to community-wide mortality during the rapid change in osmotic conditions associated with rewetting of dry soil [7, 8]. Reversal of this pattern over the following day is thus likely attributed to extensive growth across the community.

Organisms growing at the same time may be using similar resources, and changes in the organisms that are growing could be driven by changes in resources available. Specifically, cellular growth requires acquisition of carbon and nitrogen. The main sources of these elements in soil are organic, i.e., complex carbohydrates and large nitrogen molecules such as proteins, chitin and microbial cell walls. Therefore, we focused on the genetic repertoires related to these substrates and their differential abundances over time and between treatments.

Respiration of ^13^CO_2_ was produced from labeled plant-derived material, likely a mixture of plant litter and microbial bodies that consumed plant material during the previous year’s growing season. It has been previously suggested that in soils that experienced drought, microbes take up less plant derived ^13^C compounds during the dry season than in soils that didn’t experience drought [31]. Therefore, the slightly lower respiration of ^13^C in the reduced precipitation treatment suggests that microbial bodies served as a carbon source following wetup, and those were less isotopically labeled under reduced precipitation due to the lower uptake of labeled plant material. ^13^C respiration was detected throughout the week after wetup. However, this respiration was fastest in both treatments within the first 48h, implying usage of ^13^C-material by fast growers. Fast growing organisms had a smaller number of carbohydrate degrading enzymes than slow growers, implying that fast growth relies on labile carbon in addition to microbial bodies. Thus, based on genetic evidence and respiration of plant-derived C, we suggest that fast growers rely not only on labile C but also on plant-derived material for growth. However, fast growers are likely better competitors for labile carbon, thus necessitating slow growers to specialize in targeting more complex carbon sources.

Importantly, the changing pattern of carbohydrate degrading enzymes over the week-long succession implies reliance on different carbohydrates. Based on succession theory, this would be linked to the changing availability of substrates, which likely drove community composition changes. In contrast, over the same period the vast majority of growing organisms relied on the same nitrogen sources, namely ammonium and extracellular proteins. These could become available following cell death due to osmotic shock at wet-up and continued mortality during the succession. Thus, it appears that the succession we detected is driven more by carbon availability than nitrogen sources.

Our experimental strategy was designed to test for the effect of reduced precipitation in the prior growing season on microbial growth and propagation of specific functional genes following wetup. We refer to this as a legacy effect. One of the most striking findings of this study is the large magnitude of this effect, reflected in a difference in abundance of thousands of genes at t0 in organisms that grew after wetup. To gain insight into the causes of this legacy effect, we compared the growing microbial populations between precipitation treatments. Reduced precipitation resulted in less detectable growing organisms overall, although unlike previous studies [32] it did not result in lower respiration. Alphaproteobacteria response was altered by precipitation history with a delayed growth response in reduced precipitation as compared to mean annual precipitation. As field conditions were otherwise identical for both treatment groups, the explanation for the delayed response may relate to resource availability. Based on 16S rRNA qSIP, mortality after wet-up in the reduced precipitation treatment was three orders of magnitude lower than in the mean precipitation treatment. The resulting low abundance of labile carbon from cell lysis may have inhibited growth in the first two days, whereas higher mortality in the mean precipitation treatment likely supported growth of new cells. This raises the question of why lysis was less prevalent in the reduced precipitation treatment. While drying and rewetting of soil, even when they result in the same soil moisture, are very different processes [33], they both require the ability to withstand soil moisture changes. Wetup is rapid, and the organisms that survive it must be equipped to handle rapid changes in soil moisture. These same organisms would potentially be those that survived faster drying due to reduced precipitation over the prior year, hence the higher abundance at t0 of organisms that grew following wetup. We suggest that an important driver of the legacy effect was the combination of bacterial resilience to osmotic stress and the resulting decrease in mortality.

In the soils that experienced reduced precipitation, the carbohydrate degradation gene sets are different compared to gene sets in organisms growing in soils that received mean precipitation. This may point to the availability of different carbohydrates during and after dry-down between the treatment groups. Specifically, a higher fraction of the growing organisms from the reduced precipitation treatment after two days contained genes for degradation of cellulose. Thus, the capacity to degrade cellulose provided a selective advantage following wet up of soils that dried down earlier in the prior year. Further supporting this inference, increased degradation of cellulose was recently shown to be correlated to historical drought [34].

The legacy effect was surprisingly short-lived, and by the end of the week the metabolic potential of growing microorganisms that previously experienced reduced or mean precipitation converged entirely. This community “reset” could reflect convergence in resource abundance between the treatments, selecting for communities adapted to resource types historically available in this annual grassland. This reset is indicative of a resilient microbial community that converged to be similarly equipped for the next growing season. The intensity of drying/rewetting cycles is predicted to increase with climate change [35]. The experiment conducted here simulated this phenomenon; the results indicate that although the resistance of grassland soil communities to reduced precipitation is not high, their resilience likely is [36].

## Summary

Wet up, a pivotal time point in soils that experience Mediterranean climates, leads to a microbial succession process. Prior studies in Mediterranean soils revealed which organisms grow during this process, but not which traits allow them to do so. The SIP-informed metagenomic data analysis showed that growing organisms are the ones with higher fitness during dry down and have genes for polysaccharide production and export that likely provide resistance to desiccation. We report a disconnect between growth and abundance in organisms that respond to wetup. Slow growers likely rely on complex carbon sources more than fast growers, whereas fast growers are likely better competitors for labile carbon. Finally, we demonstrate a strong but short-lived legacy effect of prior water inputs, with convergence of the sets of growing organisms and their functional traits within a week after wetup.

## Methods

### Field site description

Soil samples for this study were collected from the site of a multi-year precipitation manipulation and ^13^CO_2_ labeling field experiment (sup. fig. 5)[28] at the University of California Hopland Research and Extension Center (HREC) in Northern California (39° 00’ 14.6” N, 123° 05’ 09.1” W). This region has a Mediterranean climate with cool, wet winters, and hot, dry summers. The HREC field station exists on territory originally home to the indigenous Pomo Nation, and the experiment site is an annual grassland, with naturalized annual grasses such as *Avena spp.* and *Bromus spp.* growing in soil from the Squawrock-Witherell complex. In the Fall, the soil’s pH is 7.3 with total C content of 15.1 mg/g and total N of 1.5 mg/g [28].

For two years preceding our sample collection, sixteen 3.24 m^2^ field plots dominated by *Avena spp.* were watered with either 100% (930 mm) or 50% (465 mm) of the site’s mean annual precipitation[28], and in February 2018 plants were isotopically labeled with ^13^CO_2_ for 5 days in the field during the active growing season [28, 37]. Labeled plants and associated soils remained undisturbed in the field throughout the remainder of the growing season and throughout the summer dry season. Samples for the analysis described here were collected on August 28^th^, 2018, prior to the first rainfall event of the Fall.

### Sample collection

Topsoil samples (0-15 cm, roughly 0.5 m^3^) were collected from six plots, three that had experienced mean annual precipitation and three that experienced 50% precipitation. The soils were collected before the first fall rainfall event, when average soil moisture was 3%, and transferred to the Lawrence Livermore National Laboratory where roots were removed, and soils were homogenized by sieving (2mm) to remove rocks and large plant necromass, all under ambient (hot, dry) conditions.

### ^18^O-water stable isotope probing

For SIP incubations, 5 g sieved soil from each of the field replicates (*n* = 3) was transferred to a 15 ml Nalgene flat bottom vial. One milliliter of isotopically enriched water (98 at% ^18^O-H_2_O) or natural abundance water (as a control) was slowly and evenly pipetted onto the soil and gently mixed with the pipette tip, resulting in a final average gravimetric soil moisture of 22%. After the water addition, vials were immediately sealed inside 500 ml mason jars with lids fitted with septa (to facilitate headspace sampling) and incubated at room temperature in the dark. Parallel jars were destructively harvested at multiple timepoints following the wet-up (0, 24, 48, 72, 168 h). At each harvest, vials containing soils were frozen in liquid nitrogen and then stored at −80°C. A total of 66 distinct samples were incubated.

DNA was extracted from all soil samples using a modified phenol chloroform protocol adapted from Barnard et al. (2015) [38]. Each sample was extracted in triplicate and then replicate DNA extracts were combined. For each extraction, soil (0.4 g +/- 0.001) was added to 2 ml Lysing Matrix E tube (MP Biomedicals) and extracted twice as follows: 500 μl extraction buffer (5% CTAB, 0.5 M NaCl, 240 mM K_2_HPO_4_, pH 8.0) and 500 μl 25:24:1 phenol:chloroform:isoamyl alcohol were added before shaking (FastPrep24, MP Biomedicals: 30 s, 5.5 m s^-1^). After centrifugation (16,100 x g, 5 min), residual phenol was removed using pre-spun 2 ml Phase Lock Gel tubes (5 Prime, Gaithersburg, MD, USA) with an equal volume of 24:1 chloroform:isoamyl alcohol, mixed and centrifuged (16,100 x g, 2 min). The aqueous phases from both extractions were pooled, mixed with 7 μl RNAase (10 mg/ml), mixed by inverting, and incubated at 50 °C for 10 min. 335 μL 7.5 M NH4^+^ acetate was added, mixed by inverting, incubated (4 °C, 1 h) and centrifuged (16,100 x g, 15 min). The supernatant was transferred to a new 1.7 ml tube and 1 μl Glycoblue (15 mg/ml) and 1 ml 40% PEG 6000 in 1.6 M NaCl were added, mixed by vortexing, and incubated at room temperature in the dark (2 h). After centrifugation (16,100 x g, 20 min), the pellet was rinsed with 1 ml ice-cold 70% ethanol, air-dried, resuspended in 30 μl 1xTE and stored at −80 °C.

Samples were density fractionated in a cesium chloride density gradient formed by physical density separation in an ultracentrifuge as previously described (Blazewicz et al. 2020), with minor modifications. For each sample, 5 μg of DNA in 150 μL 1xTE was mixed with 1.00 mL gradient buffer, and 4.60 mL CsCl stock (1.885 g mL^-1^) with a final average density of 1.730 g mL^-1^. Samples were loaded into 5.2 mL ultracentrifuge tubes and spun at 20 °C for 108 hours at 176,284 RCF_avg_ in a Beckman Coulter Optima XE-90 ultracentrifuge using a VTi65.2 rotor. Automated sample fractionation was performed using Lawrence Livermore National Laboratory’s high-throughput SIP pipeline ‘HT-SIP’ [27, 39, 40], which automates fractionation and clean-up tasks for the density gradient SIP protocol. Ultracentrifuge tube contents were fractionated into 36 fractions (~200 μL each) using an Agilent Technologies 1260 isocratic pump delivering water at 0.25 mL min^-1^ through a 25G needle inserted through the top of the ultracentrifuge tube. Each tube was mounted in a Beckman Coulter fraction recovery system with a side port needle inserted through the bottom. The side port needle was routed to an Agilent 1260 Infinity fraction collector. Fractions were collected in 96-well deep well plates. The density of each fraction was then measured using a Reichart AR200 digital refractometer fitted with a prism covering to facilitate measurement from 5 μL, as previously described [41]. DNA in each fraction was purified and concentrated using a Hamilton Microlab Star liquid handling system programmed to automate glycogen/PEG precipitations [40]. Washed DNA pellets were suspended in 40 μL of 1xTE and the DNA concentration of each fraction was quantified using a PicoGreen fluorescence assay. The fractions for each sample were binned into 5 groups based on density (1.6400-1.7039 g/ml, 1.7040-1.7169 g/ml, 1.7170-1.7299 g/ml, 1.7300-1.7449 g/ml, 1.450-1.7800 g/ml), and fractions within a binned group were combined and sequenced. Unfractionated DNA from each soil sample was also saved for sequencing.

### Metagenomic analyses

Biological triplicates of unfractionated DNA from the three field plots, as well as DNA from 5 density fractions per sample (*n* = 234) were sequenced using an Illumina Novaseq S4 2×150bp platform by Novogene Co. (novogene.com). Illumina adapters and phiX sequences were removed using BBmap (https://jgi.doe.gov/data-and-tools/bbtools) and the remaining reads were quality trimmed using Sickle for paired-end reads [42]. This process yielded 155·10^6^ ± 15·10^6^ (mean ± standard deviation) reads per unfractionated sample, and 38·10^6^ ± 3·10^6^ reads per density fraction.

Triplicates of unfractionated samples from each time point were co-assembled using MEGAHIT version 1.2.9 [43] with presets large-meta and minimum contig length 1000 bp. Reads from unfractionated samples were mapped to each assembly with BBwrap (requiring minimum identity 98%, https://jgi.doe.gov/data-and-tools/bbtools) and used for binning with Maxbin 2.0 [44] and MetaBAT2 [45]. Density fractions from each sample were co-assembled and binned in the same manner. Bins from each assembly were refined with the MetaWrap pipeline [46] requiring a minimum of 50% bin completion and maximum 10% bin redundancy. The resulting bins from all assemblies were dereplicated (along with bins assembled from soil samples collected from a nearby site in the winter of 2018 [21]) using dRep [47], requiring 75% completeness and under 10% contamination, yielding 503 unique metagenome assembled genomes (MAGs): 385 from the current study and 193 from the winter study. MAG taxonomy was assigned with the GTDB-tk classify workflow [48]. Three additional low-quality MAGs of Firmicutes were manually binned using ggkbase. These bins were used only for SIP analyses (not for any functional analyses), and were included because a former 16S-rRNA qSIP analysis from the same soil [8] indicated high growth rates of Firmicutes, and we were unable to automatically assemble high quality bins of this phylum.

Reads from unfractionated samples and density fractions were mapped to our MAG collection described above. Open reading frames (ORFs) were identified in MAGs with Prodigal version 2.6.3 [49] and annotated with DRAM [50]. Read counts per ORF from MAGs per sample were parsed with Rsubread [51] and then normalized and analyzed with DESeq2 [52].

### 16S rRNA gene quantification and sequencing

For 16S rRNA gene amplicon sequencing, density fractionated DNA from 330 density fractions was amplified in triplicate 10-uL reactions using primers 515 F and 806 R [53, 54]. Each reaction contained 1uL sample and 9uL of Phusion Hot Start II High Fidelity master mix (Thermo Fisher Scientific) including 1.5mM additional MgCl_2_. PCR conditions were 95 C for 2 min followed by 20 cycles of 95 C for 30 S, 64.5 C for 30 S, and 72 C for 15 S. The triplicate PCR products were then pooled and diluted 10X and used as a template in a subsequent dual indexing reaction that used the same primers including the Illumina flowcell adaptor sequences and 8-nucleotide Golay barcodes (15 cycles identical to initial amplification conditions). Resulting amplicons were purified with AMPure XP magnetic beads (Beckman Coulter) and quantified with a PicoGreen assay on a BioTek Synergy HT plate reader. Samples were pooled at equivalent concentrations, purified with the AMPure XP beads, and quantified using the KAPA Sybr Fast qPCR kit (Kapa Biosciences). Libraries were sequenced on an Illumina MiSeq instrument at Northern Arizona University’s Genetics Core Facility using a 300-cycle v2 reagent kit.

### Quantitative stable isotope probing

Atom fraction excess (AFE), growth rates, mortality rates and net rates were calculated using quantitative stable isotope probing (qSIP) [8, 55] by normalizing MAG relative abundance to the quantity of DNA per density fraction [21]. Relative abundance was calculated with coverM version 0.4.0 (https://github.com/wwood/CoverM), generating read counts (method count) for MAGs with a minimum covered fraction of 50% (method covered_fraction) and dividing by total reads per sample. Cumulative mortality was calculated using mortality rates estimated with 16S rRNA gene sequence and qPCR data as described in Blazewicz et al. 2020 [8, 56].

### CO_2_ Flux quantification

For analysis of CO_2_ and ^13^CO_2_, headspace gas samples (5 ml) were collected at each harvest with a gas-tight syringe and transferred to 20 ml wheaton serum bottles that had been purged and filled with N2 (1 atm). Total headspace CO_2_ was quantified via gas chromotography equipped with a methanizer paired with a flame ionization detector (GC2014, Shimadzu). Carbon isotopic composition (δ^13^C relative to the VPDB standard) of CO_2_ was measured with a Picarro G2201-i cavity ring down spectrometer fitted with a small sample introduction module (SSIM).

### Statistical analyses

R code used in this study is available on https://github.com/ellasiera/Wetup_traits. Cumulative mortality regression was calculated with the R base function lm based on mortality values resulting from qSIP analyses. Abundance (RPKM), read counts and breadth (fraction of MAG covered) tables of MAGs per sample were calculated with coverM (https://github.com/wwood/CoverM). For qSIP analyses, counts were converted to zero where breadth was <50%. Relative abundance was then calculated by dividing read counts per MAG by number of reads per sample. Growing organisms (*n* = 79) were defined as MAGs with detected growth for which the lower limit of the confidence interval was greater than zero. Detected organisms (*n* = 200) were those that had breadth >50% and RPKM >0 in at least one sample. Abundance boxplots of growing and non-growing organisms were plotted based on RPKM using ggplot2 [57]. Heatmaps for abundance and median atom fraction excess (AFE) were plotted using the R package heatmap3 [58]. Comparisons of gene abundance between time points and treatments were performed with DESeq2 [52] by defining the variable “condition” to include both time and treatment and comparing all conditions. P values were adjusted for multiple comparisons using FDR correction. Principal coordinates analysis was based on a Bray- Curtis dissimilarity matrix representing gene abundance per sample via the R vegan package [59], and plotted with ggplot2 [57] and ggvenn [60].

For MAG functional annotation, we used gene annotations from DRAM [50] and determined which MAGs of growing organisms contained CAZy pathways for degradation of carbohydrates in each time point. That number was divided by the number of growing organisms at each time point to get the percent of organisms with the capability to degrade specific substrates. The resulting table was plotted using ggplot2 [57]. Similarly, we analyzed and plotted inorganic and organic nitrogen cycling genes, as well as peptidases. Finally, we used the annotated gene abundances to search for patterns in genes coding for extracellular polysaccharide (EPS) biosynthesis and export, assuming these would be necessary to generate a biofilm.

## Acknowledgements

The authors thank members of the Firestone and Pett-Ridge groups who helped with field sampling and soil processing, Shufei Lei and Rohan Sachdeva for bioinformatic support, Max Li for help with the soil incubations and DNA extractions, Ryan Gini for help with GC analyses, Michaela Hayer for help with qPCR and amplicon sequencing, and Marissa Lafler for operation of the LLNL HT-SIP qSIP pipeline and the Picarro CRDS.

This research was supported by the U.S. Department of Energy (DOE), Office of Biological and Environmental Research (BER), Genomic Science Program Lawrence Livermore National Laboratory (LLNL) ‘Microbes Persist’ Soil Microbiome Scientific Focus Area SCW1632, and a subaward to UC Berkeley. Field plots, the precipitation experiment and the ^13^CO_2_ labeling study were initially generated via DOE BER awards DE-SC0020163 and DE-SC0016247 (to MKF) and awards SCW1589, and SCW1421 (to JPR); designed and maintained by Dr. Mengting Yuan, Dr.

Donald Herman, Katerina Estera-Molina, and staff of the Hopland Research and Extension Center (HREC) through the University of California Division of Agriculture and Natural Resources. Work at LLNL was performed under the auspices of the Department of Energy, Contract DE-AC52-07NA27344.

## Ethics approval and consent to participate

Not applicable

## Consent for publication

Not applicable

## Availability of data and material

All raw data used in this publication is available on the NCBI short read archive under BioProject PRJNA856348. No custom tools were used to analyze the data.

## Authors’ contributions

ETS performed all bioinformatics and statistical analyses on metagenomic data, wrote and revised the manuscript.

AG provided genomic sequences, participated in sample processing and provided input on the manuscript.

AMN assisted with graphical illustrations and provided feedback on the manuscript.

MKF advised on paper writing and analyses and provided funding for the original precipitation manipulation experiment.

JPR co-designed the experiment, provided equipment and funding, processed samples, provided feedback and revised the paper.

SJB co-designed and ran the experiment, advised on the paper, participated in writing and analyzed 16S-qSIP data.

JFB advised and participated in paper writing and analyses.

