## Supplemental figures for "Functional succession of actively growing soil microorganisms during rewetting is shaped by precipitation history"

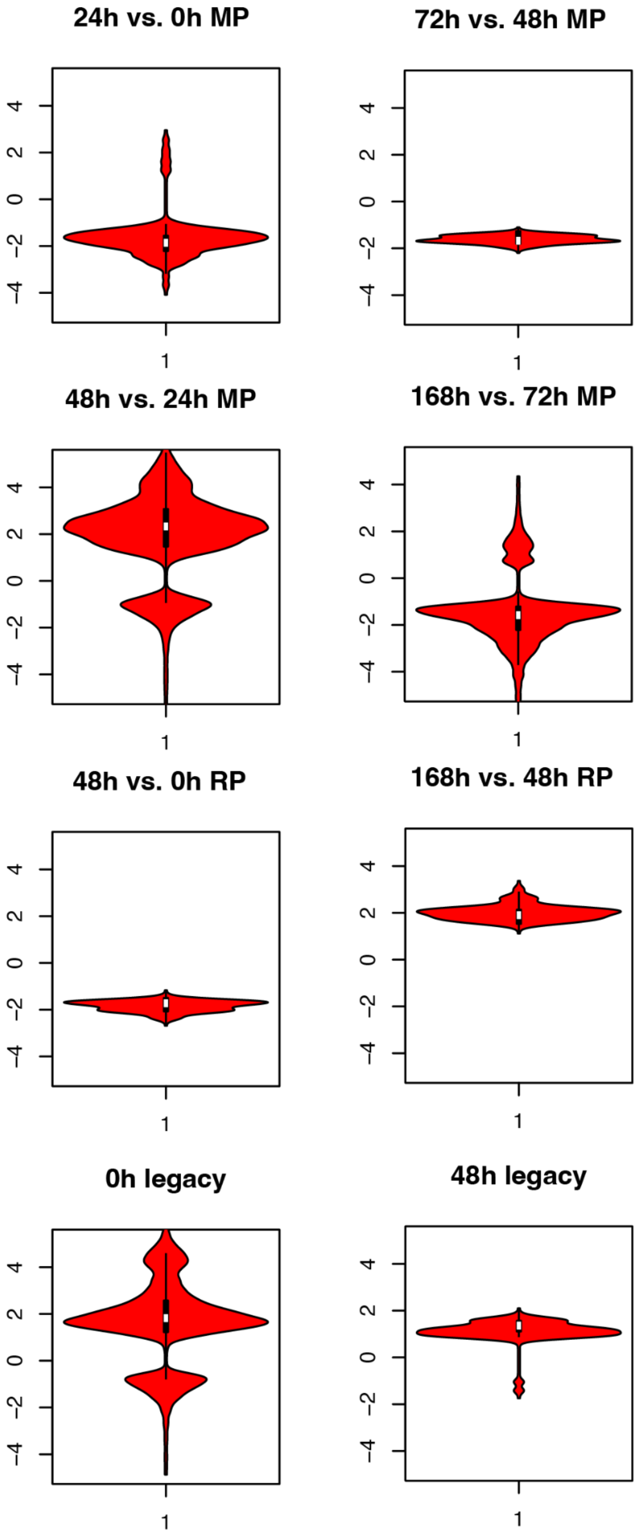


Figure 1: Distribution of significant ($p<0.05$) differential abundance of genes between time points in the mean precipitation (MP) and reduced precipitation treatments (RP), and at times 0h and 48h when comparing the treatments (legacy effect).

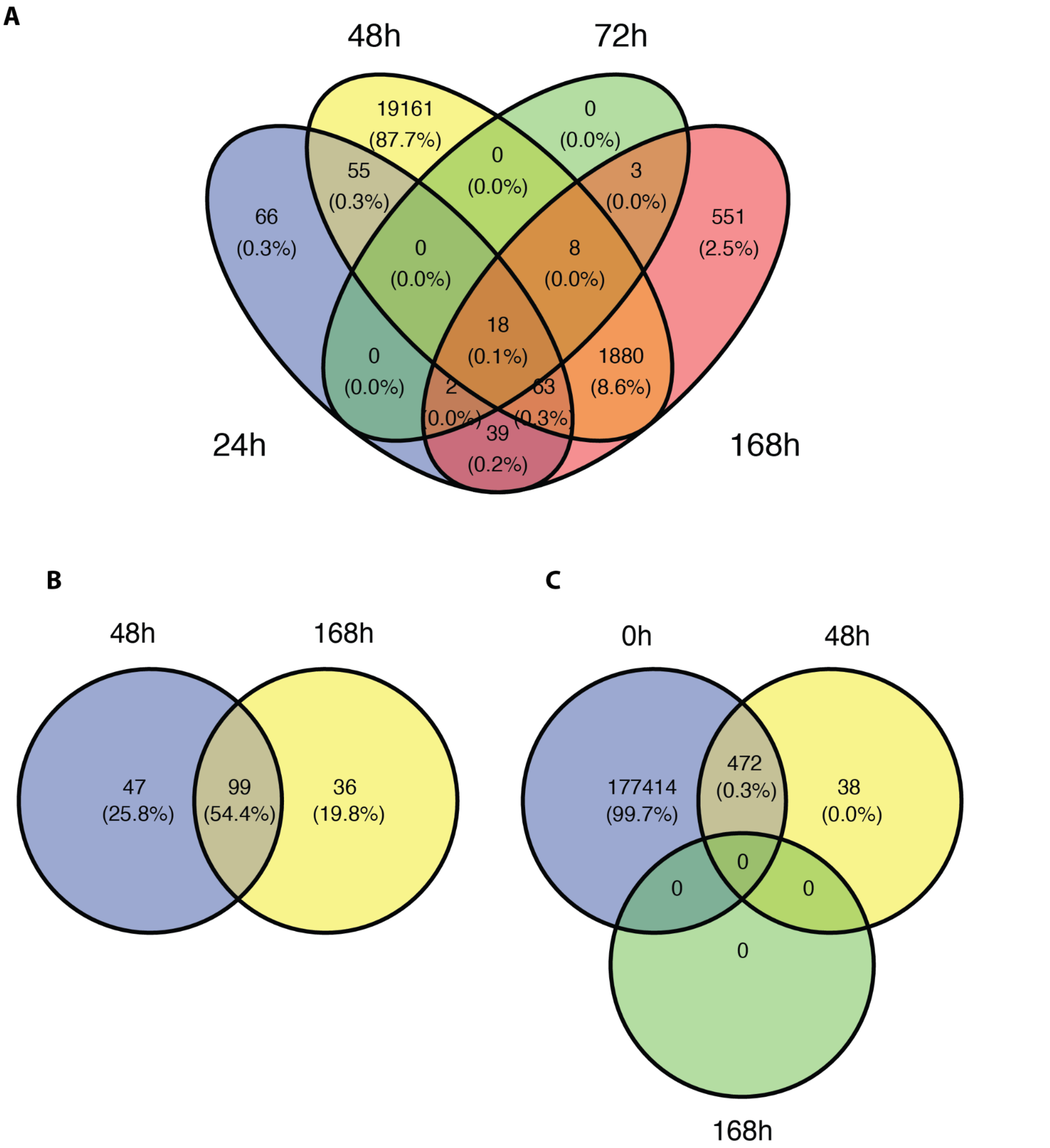


Figure 2: Shared differentially abundant genes from 77 growing organisms that were detected in at least one time point in one treatment between different time points in the (A) 100 % precipitation plots and (B) 50% precipitation plots, and (C) between comparisons of precipitation treatments (legacy effect).

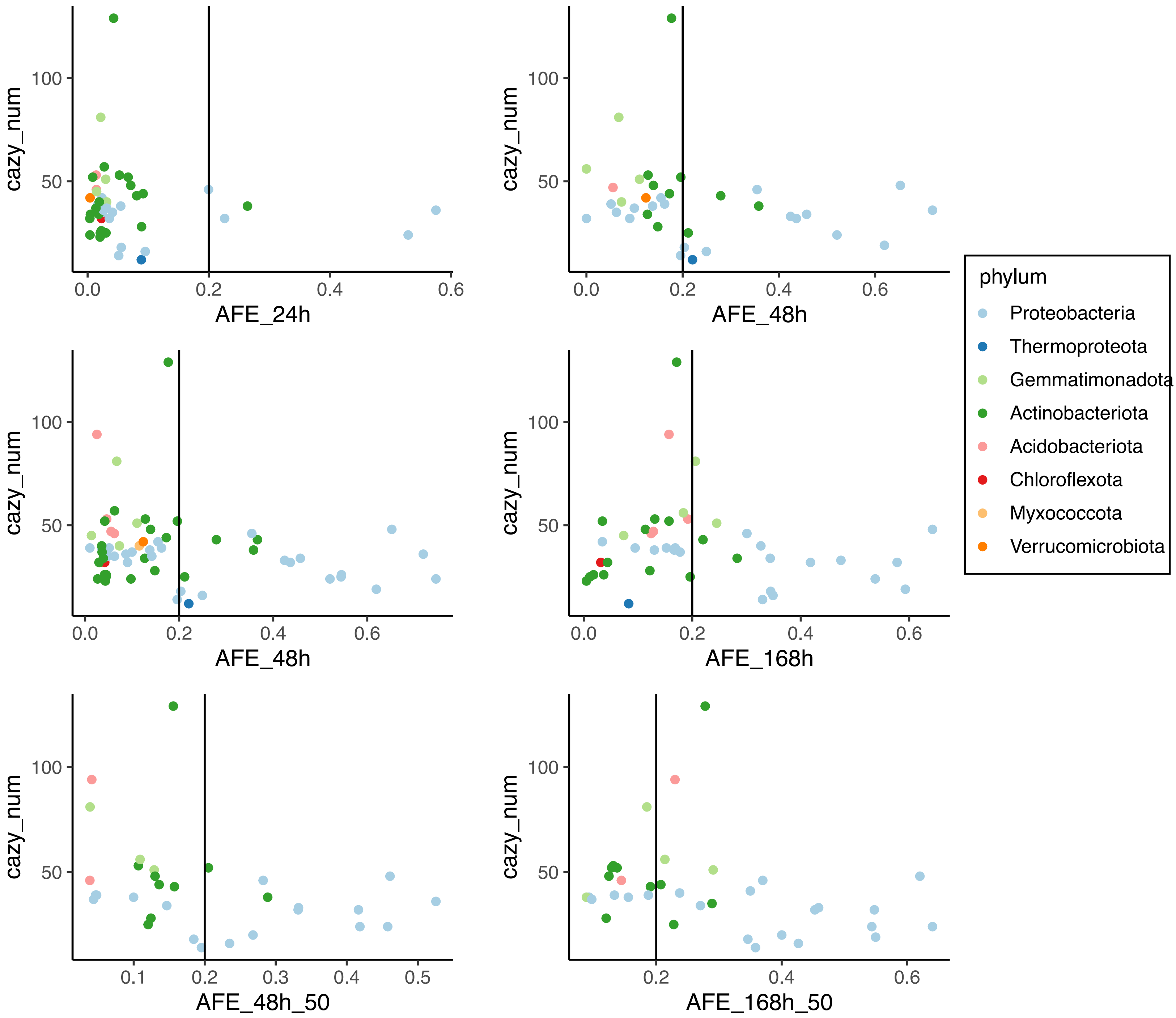


Figure 3: Number of CAZy genes in genomes of growing organisms after a soil wetup event as a function of  atom fraction excess (a proxy for growth rate). The horizontal line separates fast growing from slow growing organisms. Colors represent taxonomy. On average, slow growing organisms had 41 CAZy genes per genome as opposed to only 33 in fast growing organisms (Student’s one sided t-test, p=0.02).

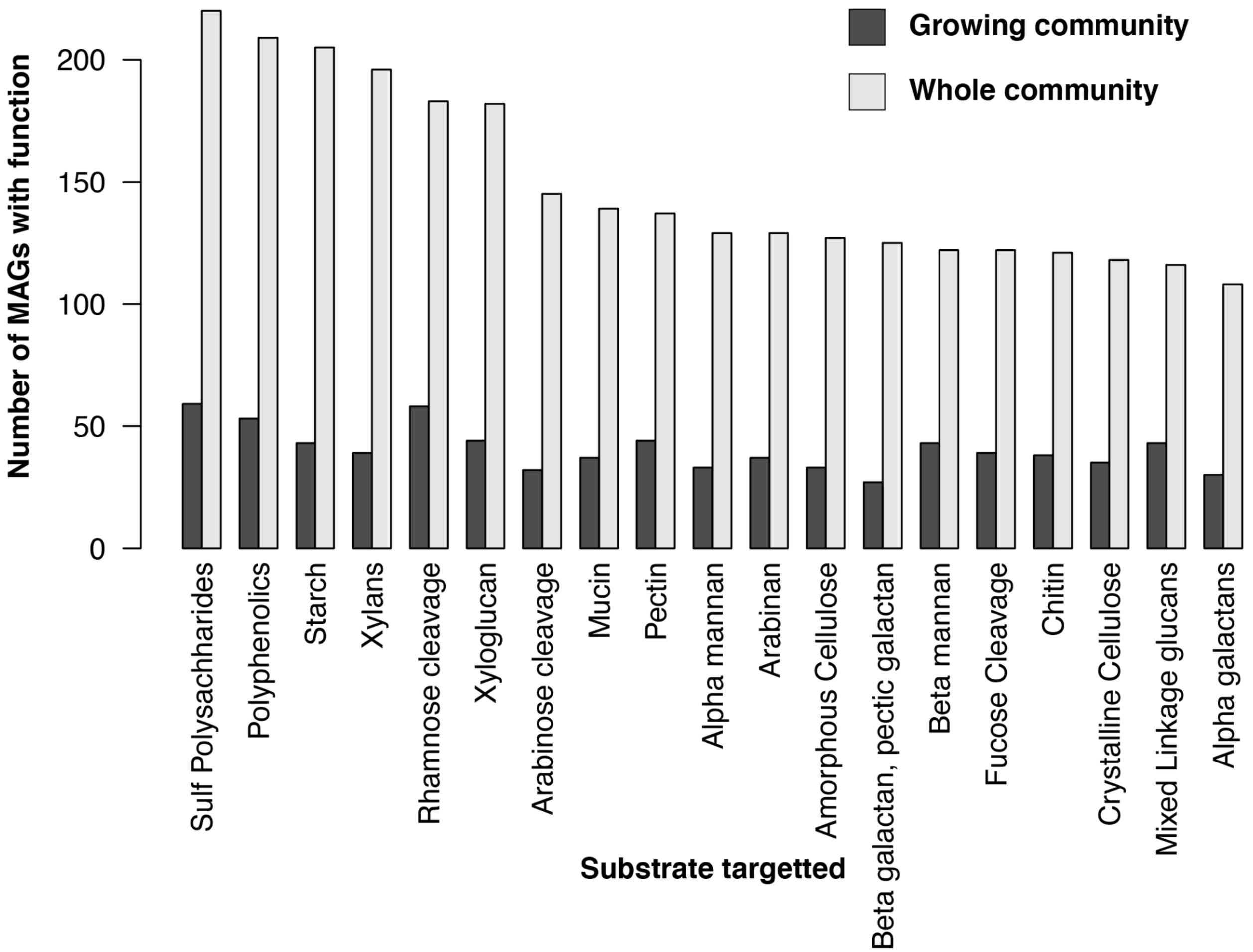


Figure 4: Counts of MAGs containing the ability to degrade specific carbohydrates in growing vs. all assembled and dereplicated MAGs ordered by counts in all MAGs. The difference in distribution of functions between the whole community and growing organisms is significantly different (Wilcoxon rank sum paired test, $p<0.001$).

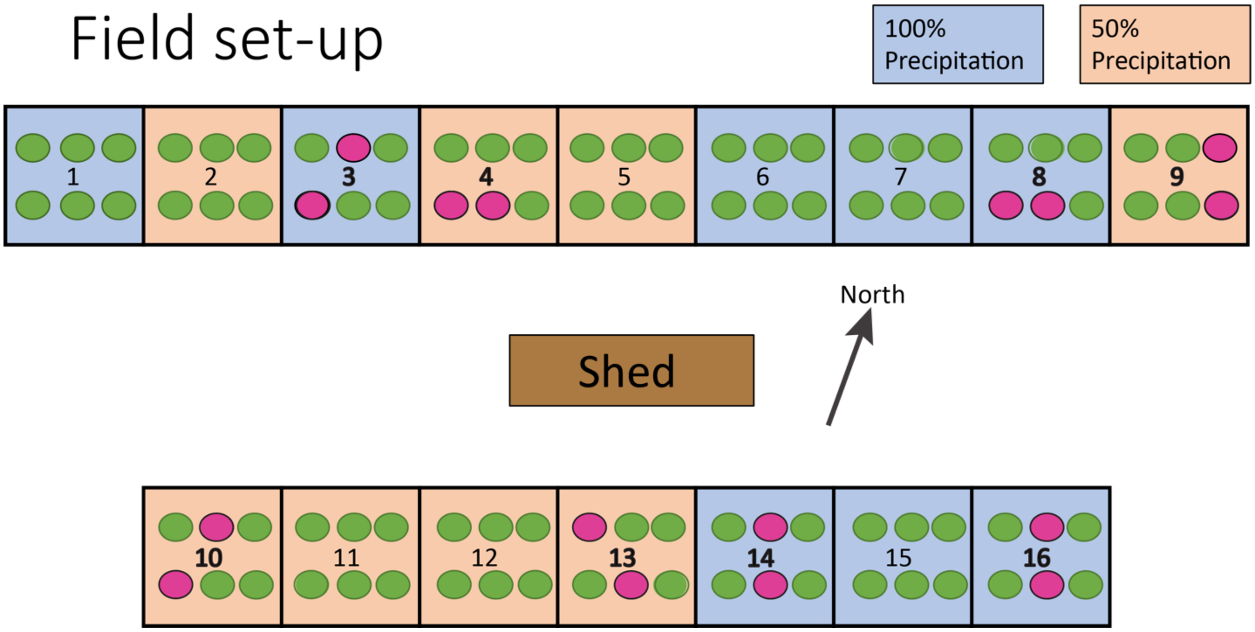


Figure 5: Field plots (1.8 * 1.8 m) in an annual grassland at the Hopland Research and Extension Center in Hopland, California. Each plot, numbered 1-16, had 6 replicate patches of *Avena* spp. surrounded by collars (20 cm radius) that were used for multiple experiments. Soil from plots 8, 9, 10, 13, 14 and 16 was collected for the wet-up experiment described here (soil was collected from collars indicated in pink, unused rings are colored green). Precipitation treatments are represented by the plot color: blue for 100% mean annual precipitation and orange for 50% mean annual precipitation.
